# Characterization of Newly Discovered Polyester Polyurethane-degrading *Methylobacterium Aquaticum* Strain A1

**DOI:** 10.1101/2024.10.18.619130

**Authors:** SeongHyeon Lee, Haemin Jeong, Injun Jung, Myounghyun Choi, Ah-Ram Kim

## Abstract

In this study, we present *Methylobacterium aquaticum* A1, a novel strain capable of degrading polyester polyurethane (PE-PUR). The attachment of *M. aquaticum* A1 to PE-PUR and its degradation capabilities were verified using Scanning Electron Microscopy (SEM) and Fourier-Transform Infrared Spectroscopy (FT-IR). Analysis of the reference genome of *M. aquaticum* revealed genes encoding enzymes with potential PE-PUR degrading activity, including esterases, lipase, proteases and amidase such as *tesA*, *pgpB*, *aes*, *aprE*, *lon*, *degQ,* and *gatA*. An esterase activity assay using *p*-nitrophenyl acetate (*p*-NPA) showed increased ester bond-cleaving activity when *M. aquaticum* A1 was exposed to polyurethane diol (PU-diol), suggesting inducible enzymatic activity involved in PE-PUR degradation. These findings highlight the potential of *M. aquaticum* A1 as a promising biocatalyst for PE-PUR degradation.

**IMPORTANCE:** Microbial biodegradation is increasingly recognized as a sustainable approach to addressing microplastic pollution. This study introduces *M. aquaticum* A1, a newly isolated strain capable of adhering to and degrading polyester polyurethane (PE-PUR), one of the most widely utilized plastics. To our knowledge, this is the first report of PE-PUR degradation by a member of the *Methylobacterium* genus. This paper provides a detailed characterization of *M. aquaticum* A1 and identifies several enzyme candidates—*tesA*, *pgpB*, *aes*, *aprE*, *lon*, *degQ,* and *gatA*—as potentially involved in the degradation process. Given that *Methylobacterium* species are known to be ecologically beneficial and inhabit diverse environments, the capacity of *M. aquaticum* A1 to degrade PE-PUR presents a promising strategy for mitigating microplastic pollution across a range of ecosystems.

---

Plastics are widely used due to their high plasticity, light weight, durability (1, 2). Over the decades, their convenience and affordability has resulted in a substantial global increase in plastic consumption. From 1950 to 2015, approximately 8,300 million tons of plastic materials were produced worldwide (3). As a result, plastic waste also increased dramatically, accumulating to 6300 million tons (3, 4). When the discarded plastic waste is exposed to UV radiation, high temperatures, wind, and other environmental stresses, it undergoes photodegradation, weathering, or fragmentation into smaller particles. These chemically degraded or physically fragmented plastics are categorized by size, with particles larger than 5 mm referred to as macroplastics, those smaller than 5 mm as microplastics, and those less than 0.1 μm as nanoplastics (5, 6, 7, 8, 9). Microplastics and nanoplastics have raised significant concerns regarding potential health risks, including oxidative stress, genotoxicity, apoptosis, necrosis, and inflammation (10, 11, 12). These plastic particles can enter the bloodstream through dietary intake of contaminated food or inhalation of airborne plastics (13, 14, 15, 16). Both microplastics and nanoplastics are persistent in the body due to their resistance to degradation. Over time, their accumulation in the bloodstream and vital organs, such as the brain, digestive system, kidney and liver, could potentially lead to long-term health problems (11, 17, 18, 19). However, effective methods for the safe management of the vast amounts of discarded plastics and their derivatives remain underdeveloped and present significant challenges (20, 21, 22).

Since the advent of plastic, the primary methods for managing plastic waste were landfilling and incineration, with the majority of plastic waste being buried in landfills and only 12% incinerated (4). However, landfills pose significant risks of soil and groundwater contamination, while incineration generates ash and gaseous toxic chemicals that contribute to both environmental and human health risks (20, 23, 24). To promote a sustainable environment, plastic recycling is indispensable (25). Currently, mechanical and chemical methods are the primary approaches used for plastic recycling, each with distinct advantages and disadvantages (21, 26). Mechanical recycling methods are advantageous due to their energy efficiency, relatively low cost, and straightforward processes, which include collection, identification, sorting, grinding, washing, agglomerating, and compounding. Most commercially used recyclates are produced through mechanical recycling (27). However, a significant drawback is that mechanical recycling heavily depends on the quality of the plastic waste that is being recycled (23, 27, 28). Chemical recycling, including thermolysis and solvolysis, can produce high-quality plastics suitable for medical applications, but it is generally more costly and involves complex processes to obtain the final products (23, 29).

Although plastic recycling will play an essential role in a sustainable future, the plastic waste problem is a complex issue that cannot be addressed through recycling alone (30, 31). Due to technical limitations in sorting plastic waste and removing contaminants, a significant portion of plastics currently remains unrecycled, ultimately ending up in landfills or being incinerated. Therefore, biodegradable plastics capable of degrading at ambient temperatures are being actively researched as complementary solutions to current recycling methods (32, 33, 34, 35). Additionally, microbial degradation of conventional plastics is being explored as another approach to complement plastic recycling (36). The most significant potential of microbial plastic degradation lies in its ability to break down microplastics (37, 38, 39). Recent studies are investigating the use of bioreactors for degrading microplastics in wastewater by utilizing specific strains of bacteria, fungi, and purified plastic-degrading enzymes, such as PETases, cutinases and lipases (40, 41, 42, 43). Microbial biodegradation offers a promising solution for addressing marine and terrestrial microplastic pollution, as conventional methods—such as landfilling, incineration, and recycling—are ineffective at removing widely dispersed microplastics (44). However, its commercial viability is currently limited by factors such as low degradation efficiency and the need to account for variables including the biochemical composition of the plastic, temperature, pH, oxygen availability, potential pathogenic risks, and overall environmental safety (23, 45, 46). So far, a few isolated microorganisms have successfully degraded polyethylene terephthalate (PET) and blended textiles (47, 48), and ongoing research is expanding to target other plastic types.

Polyurethane (PU), for example, is a versatile polymer extensively used in industries such as construction, automotive, and textiles (49, 50). PU comes in various forms depending on the type of polyol and isocyanate used in its production, with the isocyanate acting as a ‘glue’ that links the polyols together (51, 52, 53). The most common types of PU are PE-PUR (Poly<u>ester</u> Polyurethane) and PEE-PUR (Poly<u>ether</u> Polyurethane) (54). The accumulation of PU waste poses a significant environmental challenge, as its extremely slow biodegradation rate leads to prolonged persistence in the environment (55, 56). To address this issue, there is a growing interest in exploring microbial degradation as a potential solution for PU waste management (56, 57, 58). This study aimed to isolate and characterize novel microorganisms capable of degrading polyester polyurethane (PE-PUR), which is known for its high mechanical strength and chemical resistance (59, 60). By cultivating microorganisms in a minimal salts medium with PE-PUR as the sole carbon source, we identified a novel *Methylobacterium* strain capable of adhering to and degrading PE-PUR. This strain contains several enzyme candidates—*tesA*, *pgpB*, *aes*, *aprE*, *lon*, *degQ,* and *gatA*—that are potentially involved in the degradation process. Note that all PU samples used in this study are specifically PE-PUR; from this point onward, we refer to PE-PUR simply as PU.

## MATERIALS AND METHODS

### Isolation, identification, and culturing of bacteria

Environmental sample resuspended in minimal salts medium (MSM) containing 2.27g K2HPO4; 0.95g KH2PO4; 0.67g (NH4)2SO4; 2 ml Metals (6.37g Na2EDTA.2H2O; 1g ZnSO4.7H2O; 0.5g Cacl2.2H2O; 2.5g FeSO4.7H2O; 0.1g NaMoO4.2H2O; 0.1g CuSO4.5H2O; 0.2g CoCl2.6H2O; 0.52g MnSO4.H2O; 60g MgSO4.7H2O); per liter of distilled water (DW) and incubated with sterilized PU or PET film for 2 weeks for enrichment. Then, pure isolation of single colony bacteria was performed using streaking. After identification, *M. aquaticum* A1 culture was done in R2A medium (Kisan Bio^®^) containing 0.5g Casamino acid hydrolysate; 0.5g Yeast extract; 0.5g Proteose peptone; 0.5g Dextrose; 0.5g Soluble starch; 0.3g Dipotassium phosphate; 0.05g Magnesium sulfate; 0.3g Sodium pyruvate; per liter of DW or MSM, MSM + yeast extract 0.01%. For bacterial identification, 16S rRNA sequencing with BLAST was conducted by Macrogen^®^.

### Characterization of Bacterial Isolates

Phenotypic characterization of *M. aquaticum* strain A1 was done using GEN III Microplate^TM^ (Biolog^®^). The bacteria were streaked on R2A agar and incubated for 48 hours at 33°C. Inoculate a single colony into the Inoculating fluid (IF-C) until the turbidity (T) reaches 65% cell density. Pipette 100 μl into each well of the GEN III microplate and then incubate for 24-48 hours at 33°C. Absorption measurement was done at 590 nm using a microplate reader (Hidex Sense) and each well was compared quantitatively and visually with a control well.

### Scanning electron microscope (SEM)

For sample preparation, bacterial colonies grown on an R2A agar plate were cut into blocks. The agar blocks were fixed and dehydrated. First, fixation was done by submerging the blocks in 2.5% glutaraldehyde in phosphate-buffered saline (PBS) for 2 hours, room temperature (RT) (61). Then agar blocks were washed two times with PBS. Samples were dehydrated by submerging them sequentially in 30%, 50%, 70%, 80%, 95%, and 100% (vol/vol) ethanol for 10 minutes each (61). The last 100% ethanol dehydration lasted for 30 minutes. For bacteria-treated PET and PU films, samples were washed with 2% Sodium dodecyl sulfate (SDS) for 4 hours. Then it was washed three times using one-tenth diluted methanol solution (10:1, DW:Methanol) (62), and dried overnight on a clean bench. Before SEM imaging, sputter coating with platinum (Pt) was conducted. HITACHI S-4800 Scanning Electron Microscope was operated at 15.0 Kv for imaging.

### Fourier-transform infrared spectroscopy (FT-IR)

Attenuated total reflectance (ATR) and transmittance were used to detect functional groups of plastic films cultured with *M. aquaticum* A1. The measurement range was 4000-400 cm^-1^. The Varian 670/620 (Materials Characterization Lab., Unist) was used. The spectral resolution was 0.075 cm^-1^ and used an MCT detector covering a range from 6000-450 cm^-1^. The signal-to-noise ratio (5 sec): 12,000: 1 with 25% source power. ATR imaging was 1.4 μm per 1 pixel.

### Plastic film preparation

PET and PU films were cut into 1×1 cm^2^ or 2×2 cm^2^ and prepared through several steps of sterilization. Incubate plastic films in a solution of 1% (v/v) bleach and 0.7% (v/v) Tween 80 at 30°C for an hour, followed by washing with 70% ethyl alcohol (EtOH) for 30 minutes. Finally, it was washed with 30% hydrogen peroxide (H_2_O_2_) for 1 minute, rinsed with distilled water (DW), and dried overnight on a clean bench.

### Esterase activity assay

*M. aquaticum* A1 and *Escherichia coli* DH5α (*E. coli* DH5α) were grown to exponential phase in R2A and LB broth, respectively. Bacterial cells were then collected by centrifugation. After washing twice, the optical density (OD_600_) was adjusted to 0.1 with MSM. PU-diol (88 wt% in H2O, Sigma) was added at 0.01g/ml and incubated for 24 hours. Next, the supernatant was collected by centrifugation and kept on ice for the assay. 50 μl of samples were incubated with 20 mM Tris-HCl pH 7.5 and 200 μM *p*-NPA (freshly prepared in 5 mM stocks in ethanol) to a final volume of 200 μl in UV-cuvette (63). Measurement was done at 405 nm at room temperature using a UV-spectrophotometer (Shimadzu, Japan) for 2 minutes. The extinction coefficient (ε410) of *p*-NPA at pH 7.5 is 11.8 x 10^6^ cm^2^/mol, and the optical path length is 1 cm (63).

### Identification of candidate genes involved in PU degradation

To identify enzyme candidates capable of cleaving ester and urethane bonds in PU, we first employed a keyword-based search strategy using the following terms: esterase, urethanase, amidase, cutinase, protease, urease and lipase to filter relevant genes from the UniProt database. Candidates were then refined based on the following criteria:

1. EC Number Association: Enzymes with EC numbers linked to ester and urethane bond degradation (esterase: EC 3.1; protease: EC 3.4; amidase: 3.5) were prioritized.
2. Gene Annotation: Candidates with experimentally validated gene annotations were selected to ensure reliability.
3. Motif matching: Candidates containing the following motifs—hydrolase (GGGX), serine hydrolase (GXSXG), and catalytic triads (Ser-Asp-His/Glu or Ser-cisSer-Lys)—were selected for their potential roles in ester and urethane bond cleavage. To evaluate catalytic triads as structural motifs, AlphaFold3 predictions are employed.

In the reference genome of *Methylobacterium aquaticum* MA-22A, the following candidate genes were identified as potential PU-degrading enzyme genes: *tesA*, *pgpB*, *aes*, *aprE*, *lon*, *degQ*, and *gatA*.

## RESULTS

### Identification of a novel strain *M. aquaticum* A1

A bacterial strain was isolated from a minimal salts medium (MSM) culture containing PU film during the screening process. The colonies exhibited characteristic pink to red pigmentation. The cells are rod-shaped, measuring 2.5 to 5 μm in length, and are classified as Gram-negative (**Fig. 1A**). 16S rRNA sequencing confirmed the strain’s identity as *Methylobacterium aquaticum*, which we have designated *M. aquaticum* A1. Phylogenetic tree analysis revealed that *M. aquaticum* A1 exhibited a distinction from two other *M. aquaticum* species listed in the public database (**Fig. 1B**). Its phenotypic characteristics were analyzed using a GEN III Microplate^TM^ (Biolog^®^) to identify differences from the type strain **(Table. 1)**. The carbon source utilization assay results are as follows: *M. aquaticum* A1 can utilize Dextrin, Gentiobiose, D-melibiose, D-fructose, D-galactose, 3-Methyl glucose, D-fucose, L-fucose, L-rhamnose, D-glucose-6-PO4, D-fructose-6-PO4, D-aspartic acid, L-histidine, L-pyroglutamic acid, D-galacturonic acid, L-galactonic acid lactone, D-gluconic acid, D-glucuronic acid, Glucuronamide, Mucic acid, D-saccharic acid, *p*-hydroxy-phenylacetic acid, D-lactic acid methyl ester, L-lactic acid, Citric acid, a-keto-glutaric acid, D-malic acid, Tween 40, a-hydroxy-butyric acid, β-hydroxy-D, L-butyric acid, a-keto-butyric acid, Acetoacetic acid, Propionic acid, Acetic acid, Formic acid. And cannot utilize D-maltose, D-trehalose, D-cellobiose, Sucrose, D-turanose, Stachyose, D-raffinose, a-D-lactose, b-methyl-D-glucoside, D-salicin, N-acetyl-D-glucosamine, N-acetyl-D-mannosamine, N-acetyl-D-galactosamine, N-acetyl neuraminic acid, a-D-glucose, D-mannose, Inosine, D-sorbitol, D-mannitol, D-arabitol, myo-inositol, Glycerol, D-serine, Gelatin, Glycyl-L-proline, L-alanine, L-arginine, L-aspartic acid, L-glutamic acid, L-serine, Pectin, Quinic acid, Methyl pyruvate, L-malic acid, Bromo-succinic acid, r-amino-butyric acid. In the utilization assay, it showed 80% similarity with the type strain *M. aquaticum* GR16 (64).

The chemical sensitivity assay results are as follows: It can grow at pH 6, 1% NaCl, 1% sodium lactate, Rifamycin SV, Minocycline, Lincomycin, Vancomycin, Tetrazolium violet, Tetrazolium blue, Nalidixic acid, Potassium tellurite, Aztreonam. And cannot grow on pH 5, 4% NaCl, Fusidic acid, D-serine, Macrolide, Guanidine HCl, Niaproof 4, Lithium chloride, Sodium butyrate, Sodium bromate.

**FIG 1.**
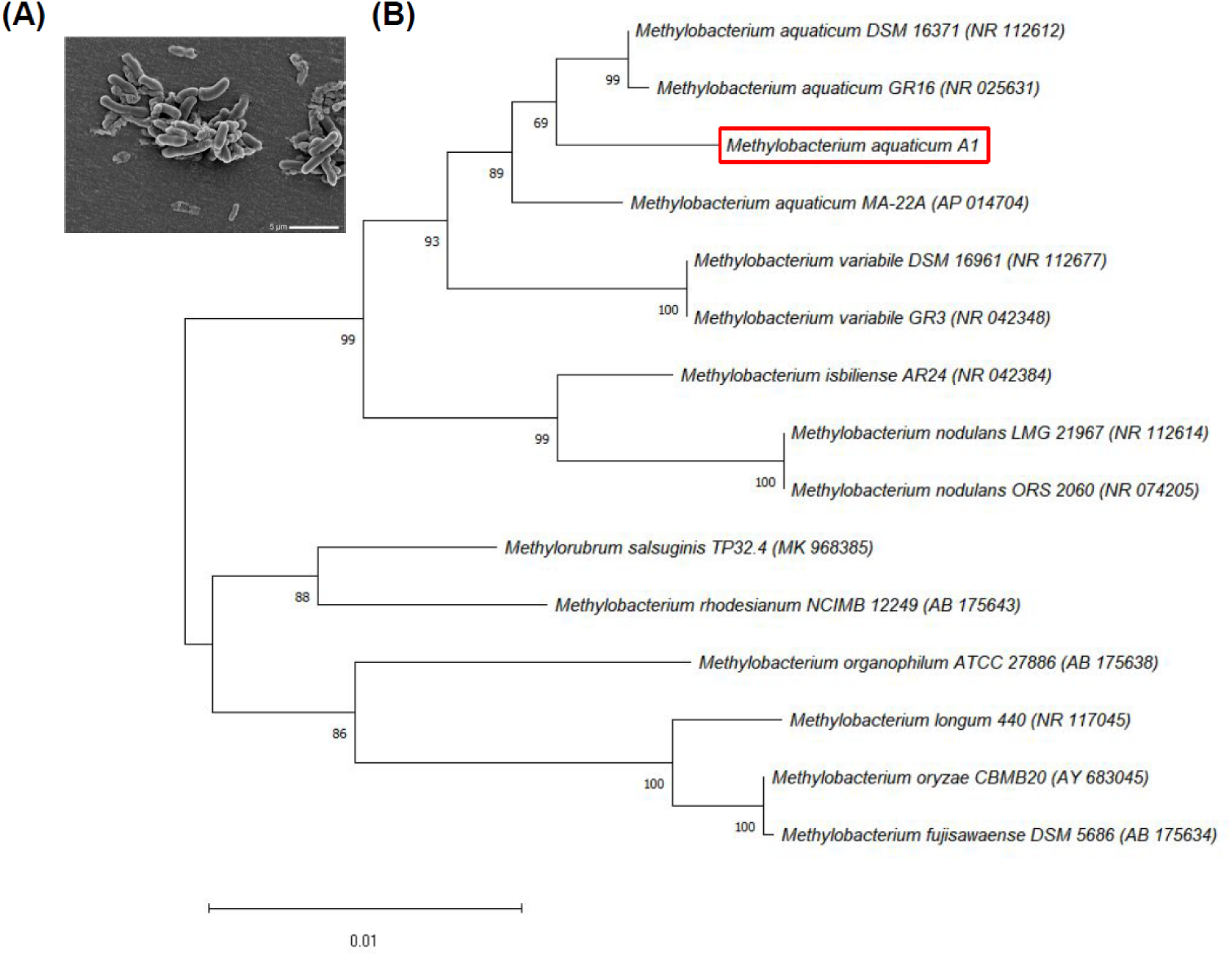
Phylogenetic analysis of newly discovered *M. aquaticum* A1 was based on its 16S rRNA sequence (1,323 bps). (A) SEM image of *M. aquaticum* A1 colony after 7 days of incubation on R2A agar at 30°C. (B) The phylogenetic tree was constructed by Molecular Evolutionary Genetics Analysis 11 (MEGA11) using the Neighbor-joining method. Bootstrap tests were performed with 1,000 replicates. The scale bar represents sequence divergence, and the bracket indicates the NCBI accession number.

**Table 1.**
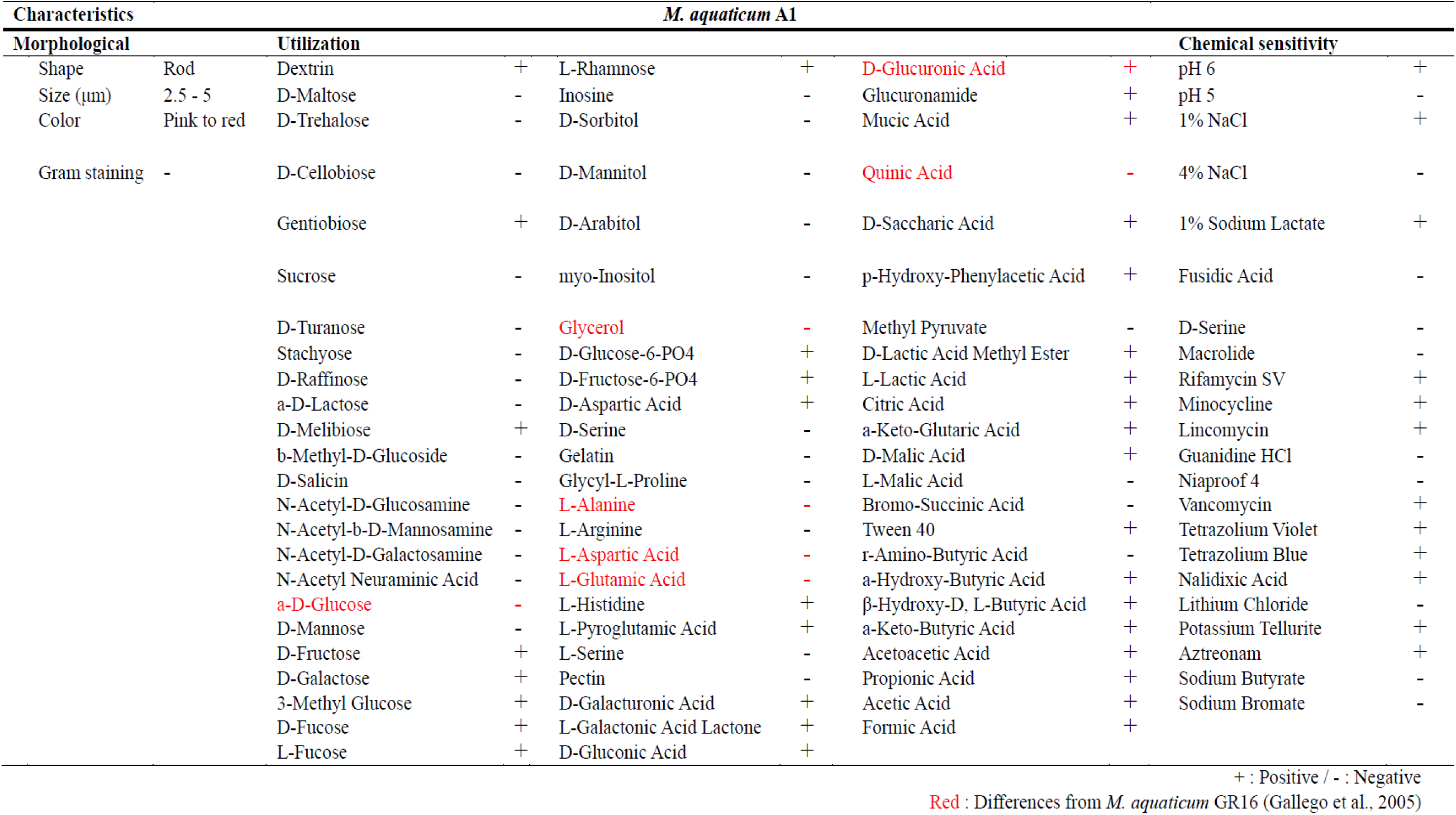
Characterization of isolated strain Al. The phenotypic characteristics of *M. aquaticum* A1, including the utilization of carbon sources and chemical sensitivity, were analyzed using the GEN III microplate^TM^ (Biolog^®^).

### *M. aquaticum* A1 formed biofilm on PU and PET films

Bacterial adherence is crucial for plastic degradation (65). To examine the attachment of *M. aquaticum* A1 to PU film, the strain was incubated for 7 days in MSM supplemented with 0.01% yeast extract and R2A media, respectively, along with PU film. Visual inspection confirmed that *M. aquaticum* A1 formed biofilms on PU film in both MSM supplemented with 0.01% yeast extract and R2A media **(Fig. S1A and Fig S1B)**. The 0.01% yeast extract added to MSM provides minimal nutrients to support early biofilm formation by the inoculated microorganisms, whereas R2A media supplied sufficient nutrients for overall growth. Biofilms were formed in both media, but they were thicker and more extensive when grown in R2A media. For cellular-level observation, the incubated PU film was washed, fixed, dehydrated, and Pt-coated for SEM analysis. SEM images showed pili-like structures for attachment and the localized formation of substances, likely to be extracellular polymeric substances between bacteria (**Fig. 2A**). These features were not seen in SEM images of cultures grown on agar plates (**Fig. 1A**). Notably, a similar level of biofilm formation was observed on PET film, another widely used synthetic polymer, in R2A medium (**Fig. 2B and Fig. S1D)**. However, in MSM with 0.01% yeast extract, visual inspection revealed reduced biofilm formation on PET film compared to PU film **(Fig. S1C)**, suggesting a tendency for biofilm formation on various plastic surfaces rather than exclusively on PU, although biofilm formation on PET film is less effective than on PU film in the MSM-based medium.

**FIG 2.**
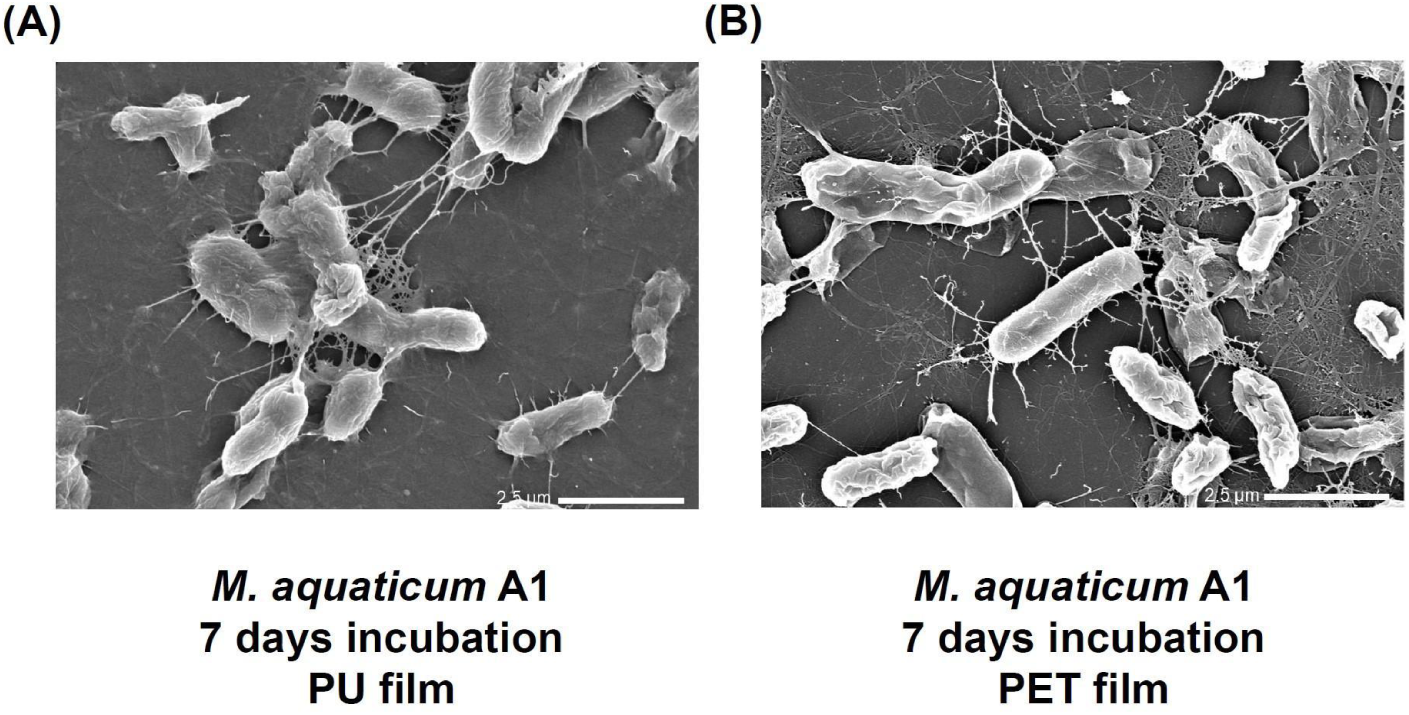
Adherence of *M. aquaticum* A1 to PU and PET films was observed using SEM analysis. PU and PET films were incubated with *M. aquaticum* A1 in R2A media at 30°C. (A) PU film incubated with *M. aquaticum* A1 for 7 days. (B) PET film incubated with *M. aquaticum* A1 for 7 days.

### The surface erosion of the PU film by *M. aquaticum* A1 was observed

To assess plastic degradation in regions with biofilm formation, PU and PET films were incubated with *M. aquaticum* A1 for 30 days. Sterilized PU and PET films were incubated in MSM supplemented with 0.01% yeast extract at 30°C. *M. aquaticum* A1 was inoculated at an initial OD_600_ of 0.1. After 30 days, the films were washed with sodium dodecyl sulfate (SDS) and methanol to remove bacterial cells, followed by SEM analysis. Compared to the negative control (**Fig. 3A**), the PET film incubated with *M. aquaticum* A1 for 30 days showed no signs of erosion (**Fig. 3C**), despite biofilm formation. In contrast, clear signs of erosion were observed on the surface of the PU film (**Fig 3B**), confirming the ability of *M. aquaticum* A1 to degrade PU. The dark areas in the figure 3B represent recessed regions.

**FIG 3.**
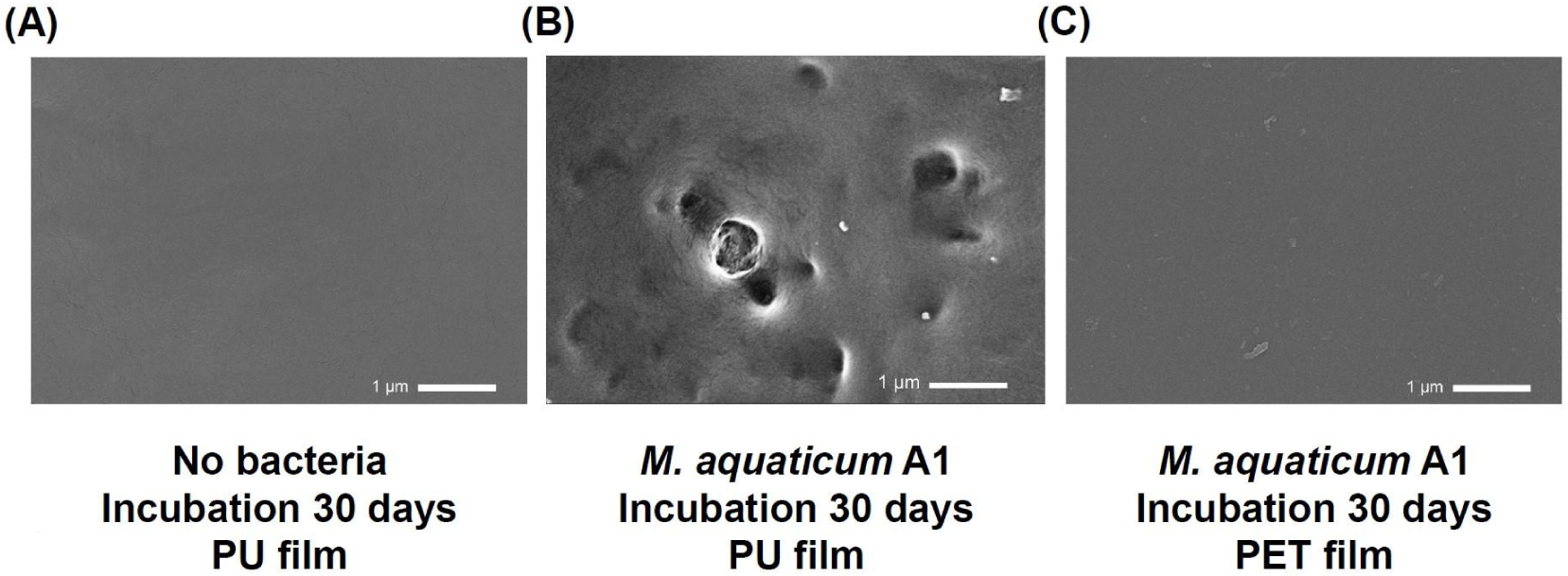
Surface recessions of the PU film were caused by *M. aquaticum* A1. (A) No bacteria incubation with PU film for 30 days. (B) *M. aquaticum* A1 incubation with PU film for 30 days. (C) *M. aquaticum* A1 incubation with PET film for 30 days. Sterilized PU and PET films, along with *M. aquaticum* A1, were incubated in MSM with 0.01% yeast extract at 30°C.

### FT-IR confirmed the PU degradation by *M. aquaticum* A1

In the PU polymer, the carbonyl (C=O) group is characteristic of both ester bonds (-R-C(=O)-O-R’) and urethane linkages (-NH-C(=O)-O-C-) (66). To assess whether the observed erosion of the PU film was due to bacteria-mediated hydrolysis of ester bonds and/or urethane linkages, the chemical composition of the PU film surface was analyzed using Fourier Transform Infrared (FT-IR) spectroscopy, which identifies chemical compounds and elucidates molecular structures based on IR light absorption (67, 68). FT-IR analysis showed a reduction in the C=O stretching band for ester bonds at around 1730 cm^-1^ (**Fig. 4B: Panel 3)**, along with a decrease in the peak corresponding to C-O stretching at 1222-1076 cm^-1^ (**Fig. 4B: Panel 1-2)**, indicating the cleavage of ester bonds within the PU film (69, 70, 71). Additionally, the reduction in the peak at approximately 1704 cm^-1^ (**Fig. 4B: Panel 3)**, representing the C=O stretching of urethane bonds (72), along with the appearance of a new peak in the 3300-3500 cm^-1^ region (**Fig. 4C: Inset)** corresponding to free N-H stretching (73, 74), indicates the formation of amine groups (-NH_2_), a byproduct of urethane bond (-NH-C(=O)-O-C-) breakage (72, 73). The presence of the amine signal implies the possibility of urethane linkage hydrolysis. In contrast, other functional group peaks unrelated to ester bonds and urethane linkages showed no significant changes (**Fig. 4A**). These findings suggest that the observed degradation was due to bacteria-mediated cleavage of both ester bonds and urethane linkages.

**FIG 4.**
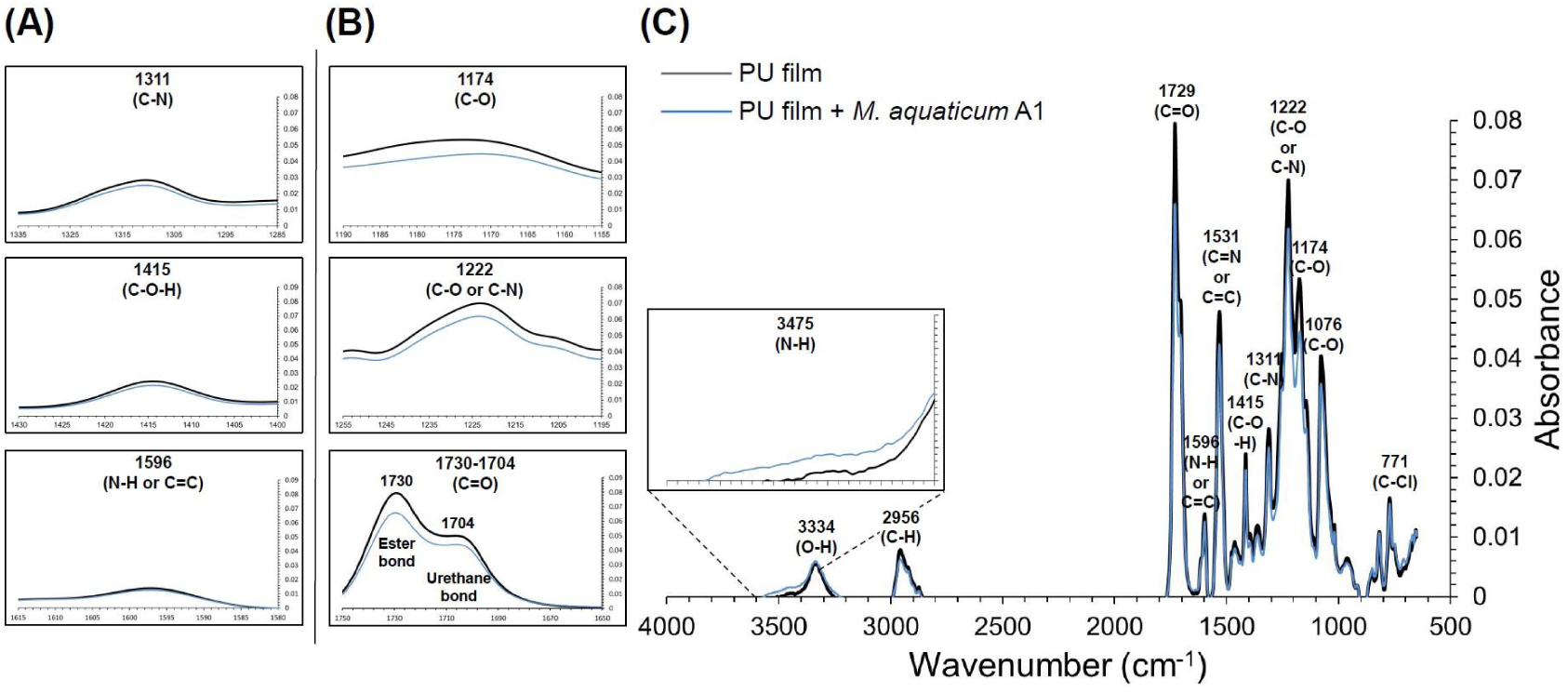
*M. aquaticum* strain A1 reduced the carbonyl group of the PU film. PU film (No bacteria), PU film + MA1 (incubated with *M. aquaticum* A1). (A) Unchanged peaks; amines (C-N, 1311 cm^-1^); (C-O-H, 1415 cm^-1^); primary amines, secondary amines, primary amides, secondary amides or alkyne (stretch) (N-H or C=C, 1596 cm^-1^). (B) Changed peaks; esters (C-O, 1174 cm^-1^); amines (C-O or C-N, 1222 cm^-1^); carbonyl groups (C=O, 1730-1704 cm^-1^). (C) FT-IR spectrum of *M. aquaticum* A1-incubated PU film in the range of 4000-400 cm^-1^. Inset: N-H stretching at 3475 cm^-1^.

### Enhanced ester bond-cleaving activity of *M. aquaticum* A1 in the presence of PU

To further confirm the involvement of enzyme-mediated ester bond hydrolysis in PU degradation, ester bond-cleaving activity was evaluated in *M. aquaticum* A1 and the negative control, *E. coli*, both in the presence and absence of PU-diol (a soluble PU monomer). The activity was quantified by measuring the production of *p*-nitrophenol, which absorbs light at 410 nm (75, 76). *p*-Nitrophenol is formed as a byproduct during the hydrolysis of ester bond-containing *p*-nitrophenyl acetate (*p*-NPA), a reaction catalyzed by esterases and other ester bond-cleaving enzymes (77). When PU-diol was added to MSM, the ester bond-cleaving activity of *M. aquaticum* A1 significantly increased compared to that without PU-diol (**Fig. 5**). In contrast, no significant difference in ester bond-cleaving activity was observed in the *E. coli* treatment groups, suggesting that *M. aquaticum* A1 enhances ester bond cleavage enzyme activity during PU degradation under carbon source-limited conditions.

**FIG 5.**
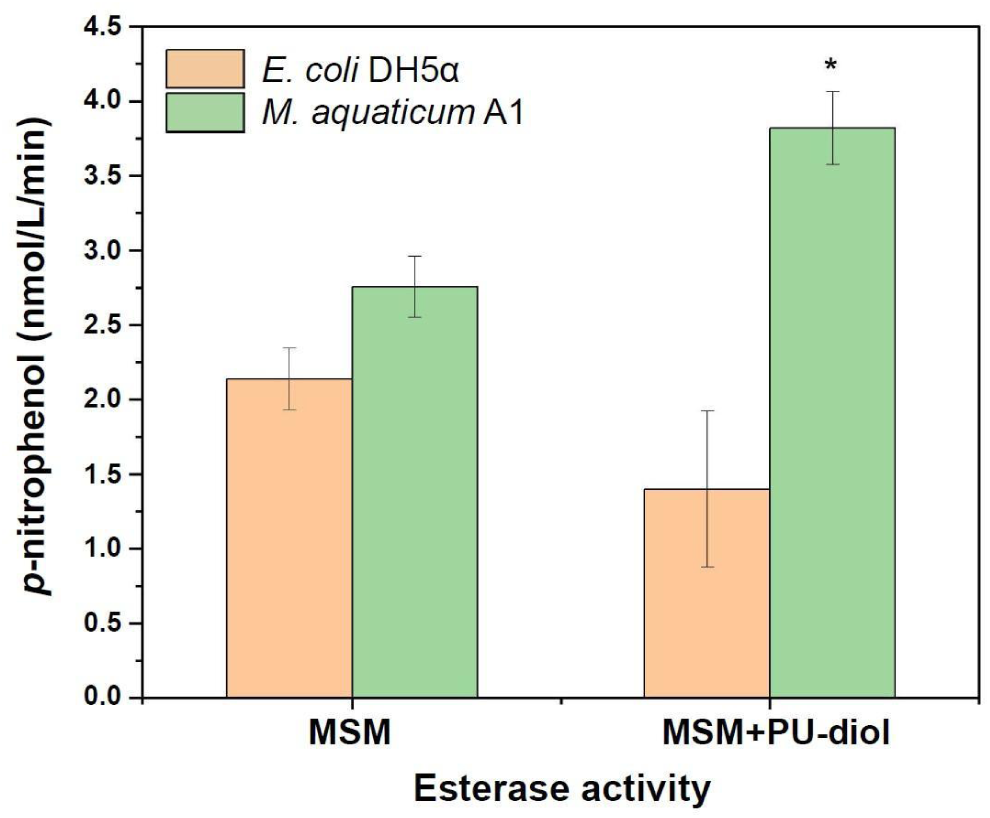
Ester bond-cleaving activity of *M. aquaticum* A1 was assessed with and without PU-diol. *M. aquaticum* A1 and *E. coli* DH5α (negative control) were first cultured overnight in R2A and LB media, respectively, and then inoculated into MSM or MSM + PU-diol at an OD of 0.1. The experiments were conducted in triplicate, with each divided into two technical replicates for measurement (\**p*-value < 0.05).

### Reference genome of *M. aquaticum* encodes potential enzymes for ester and urethane bond cleavage

We searched for *M. aquaticum* genes encoding enzymes likely to catalyze PU degradation. Various bacterial enzymes, including esterases, urethanase, amidase, cutinase, protease, lipase, and urease have been proposed to cleave ester or urethane bonds at different sites within PU (78, 79, 80, 81, 82). Among these, esterase and protease, and amidase are considered one of the most promising candidates due to its demonstrated moderate degradation efficiency in cleaving ester and urethane bonds, respectively, in PU (83, 84). Genomic analysis of the reference strain *Methylobacterium aquaticum* MA-22A revealed the presence of esterase and lipase genes, including *tesA*, *pgpB*, and *aes*, each containing either a hydrolase motif (GGGX) or a serine hydrolase motif (GXSXG) **(Fig. S2)**, both of which are known for catalyzing ester bond hydrolysis (85). Additionally, the *M. aquaticum* reference genome contains protease genes such as *aprE*, *lon*, and *degQ*, each featuring not only serine hydrolase motifs but also a catalytic triad (Ser-Asp-His/Glu), suggesting potential activity in both peptide bond and urethane bond cleavage (84, 85, 86, 87, 88). Furthermore, an amidase gene was identified with a highly conserved motif (GGSS) and a catalytic triad (Ser-cisSer-Lys), which has recently been associated with urethane bond cleavage (89).

## CONCLUSION

In this study, we identified a novel *Methylobacterium aquaticum* strain capable of degrading polyester polyurethane, as confirmed by SEM and FT-IR analysis. A notable increase in the ester bond-cleaving activity of *M. aquaticum* A1 in the presence of PU-diol under carbon source-limited conditions suggests inducible enzymatic activity, a molecular mechanism contributing to polyester polyurethane degradation. Several esterase, lipase, protease and amidase genes that may be involved in breaking ester or urethane bonds were identified. This study provides the first evidence of polyester polyurethane degradation by *M. aquaticum*.

## DISCUSSION

The *Methylobacterium* genus are facultative methylotrophs, primarily consuming methanol and other single-carbon molecules as a carbon source (90). The discovery that *Methylobacterium aquaticum* A1 can degrade not only single-carbon molecules but also the synthetic carbon-based polyester polyurethane highlights the metabolic flexibility of methanotrophs. This finding suggests that these organisms can adapt their pathways to break down complex carbon-based molecules when preferred single-carbon substrates, such as methane or methanol, are limited. Previous studies have shown that certain *Methylobacterium* species can degrade complex organic compounds under specific conditions; for instance, *Methylocella silvestris* can oxidize both methane and multi-carbon compounds like propane and butane (91), while *Methylocystis parvus* has demonstrated potential to degrade polyhydroxybutyrate (PHB), a biodegradable plastic (92, 93). However, the ability of methanotrophs to degrade PU polymers has not been previously documented. This result expands our understanding of methanotroph ecology, suggesting that they may play a broader role in carbon cycling than previously recognized. *Methylobacterium aquaticum* is an ecologically beneficial bacterium commonly found in natural environments such as water, soil, and plant surfaces (64, 94). These bacteria colonize plant surfaces, where they utilize methanol released during plant growth, providing various benefits to the host plant. For example, they produce hormones like auxin and cytokinin, which promotes plant growth and strengthens root systems, and accumulate β-carotene within their cells, helping to mitigate UV stress for both the bacteria and their host plants (95, 96, 97, 98, 99, 100, 101, 102). Additionally, *M. aquaticum* decomposes methane and methanol in soil and aquatic environments, producing beneficial byproducts and acting as a hidden regulator within ecosystems (103, 104). *Methylobacterium aquaticum* also exhibits high survivability in diverse conditions due to its ability to utilize various carbon sources, its high UV resistance, and its moderate tolerance to temperature, pH, and desiccation (105). Ecologically beneficial characteristics, combined with its high survivability positions the PU-degrading methanotrophs as potentially valuable agents for bioremediation, particularly in the context of microplastic management.

While this study demonstrated that the *M. aquaticum* strain is capable of degrading PU and investigated its potential degradation mechanisms, several limitations remain. First, the microbial PU degradation efficiency is relatively low. The slight surface erosion observed under electron microscopy after 30 days of incubation indicates that enhancing this PU-degrading capability is necessary for potential commercial applications. Future work could employ serial passage, also known as adaptive evolution, by applying selective pressure in media where PU is the primary carbon source while restricting other major carbon sources, to obtain *M. aquaticum* variants with improved PU degradation ability.

Secondly, while FT-IR is highly effective for identifying chemical changes and tracking specific functional groups, it has limitations in providing quantitative data on the exact amounts of by-products formed during PU degradation, such as diethylene glycol, butanediol, and trimethylolpropane from ester bond cleavage (106, 107), as well as polyols and amines from urethane bond cleavage (108, 109). To quantify these by-products and accurately determine the extent of PU degradation, techniques such as gas chromatography–mass spectrometry (GC–MS) or high-performance liquid chromatography–mass spectrometry (HPLC-MS) should be employed.

Lastly, although we identified a group of candidate enzymes—*tesA*, *pgpB*, *aes*, *aprE*, *lon*, *degQ,* and *gatA*—potentially involved in ester bond and urethane linkage hydrolysis in PU, we were unable to pinpoint the specific enzymes directly responsible for PU degradation. For example, while the *aes* gene contains the same serine hydrolase motif as those found in known PU-degrading enzymes (85), this hypothesis requires empirical confirmation. Substrate specificity depends on more than just the presence of specific functional motifs. The precisely arranged active site architecture of the enzyme, along with its dynamics within the protein, plays a crucial role in specifically binding and cleaving urethane bonds (110). To identify the PU-degrading enzymes, specific enzyme inhibitors can be added to a minimal salt medium containing Impranil^®^ (a water-dispersible PU substrate with urethane linkages). By monitoring microbial growth under these conditions, specific enzymes involved in PU degradation can be identified. Additionally, protein purification combined with mass spectrometry could be employed to isolate and identify active enzymes that degrade PU substrates such as Impranil^®^. Alternatively, targeted gene knockouts using CRISPR-Cas9 or its variants could systematically disable the candidate enzymes, followed by assessments of their impact on PU degradation.

## ACKNOWLEDGMENTS

We thank Jonghwan Yoon and Byumjune Park for their valuable assistance in predicting and visualizing the 3D structures of candidate enzymes, as well as analyzing their catalytic sites.

## REFERENCES

1. Sangroniz, A., Zhu, J. B., Tang, X., Etxeberria, A., Chen, E. Y. X., & Sardon, H. (2019). Packaging materials with desired mechanical and barrier properties and full chemical recyclability. Nature Communications, 10(1), 3559.

2. Joel, F. R. (1995). Polymer science & technology: introduction to polymer science. Eds, 3, 4–9.

3. Beydoun, K., & Klankermayer, J. (2020). Efficient plastic waste recycling to value added products by integrated biomass processing. ChemSusChem, 13(3), 488 492.

4. Geyer, R., Jambeck, J. R., & Law, K. L. (2017). Production, use, and fate of all plastics ever made. Science advances, 3(7), e1700782.

5. Masry, M., Rossignol, S., Gardette, J. L., Therias, S., Bussière, P. O., & Wong-Wah-Chung, P. (2021). Characteristics, fate, and impact of marine plastic debris exposed to sunlight: A review. Marine Pollution Bulletin, 171, 112701.

6. Cai, L., Wang, J., Peng, J., Wu, Z., & Tan, X. (2018). Observation of the degradation of three types of plastic pellets exposed to UV irradiation in three different environments. Science of the Total Environment, 628, 740–747.

7. Corcoran, P. L., Biesinger, M. C., & Grifi, M. (2009). Plastics and beaches: a degrading relationship. Marine pollution bulletin, 58(1), 80–84.

8. Danso, D., Chow, J., & Streit, W. R. (2019). Plastics: environmental and biotechnological perspectives on microbial degradation. Applied and environmental microbiology, 85(19), e01095 19.

9. Koelmans, A. A., Redondo-Hasselerharm, P. E., Nor, N. H. M., de Ruijter, V. N., Mintenig, S. M., & Kooi, M. (2022). Risk assessment of microplastic particles. Nature Reviews Materials, 7(2), 138–152.

10. Hu, M., & Palić, D. (2020). Micro-and nano-plastics activation of oxidative and inflammatory adverse outcome pathways. Redox biology, 37, 101620.

11. Feng, Y., Tu, C., Li, R., Wu, D., Yang, J., Xia, Y., … & Luo, Y. (2023). A systematic review of the impacts of exposure to micro-and nano-plastics on human tissue accumulation and health. Eco-Environment & Health.

12. Van Tienhoven, E. A. E., Korbee, D., Schipper, L., Verharen, H. W., & De Jong, W. H. (2006). In vitro and in vivo (cyto) toxicity assays using PVC and LDPE as model materials. Journal of Biomedical Materials Research Part A: An Official Journal of The Society for Biomaterials, The Japanese Society for Biomaterials, and The Australian Society for Biomaterials and the Korean Society for Biomaterials, 78(1), 175–182.

13. Wright, S. L., & Kelly, F. J. (2017). Plastic and human health: a micro issue?. Environmental science & technology, 51(12), 6634–6647.

14. Deng, Y., Zhang, Y., Lemos, B., & Ren, H. (2017). Tissue accumulation of microplastics in mice and biomarker responses suggest widespread health risks of exposure. Scientific reports, 7(1), 46687.

15. Waring, R. H., Harris, R. M., & Mitchell, S. C. (2018). Plastic contamination of the food chain: A threat to human health?. Maturitas, 115, 64–68.

16. Prata, J. C., da Costa, J. P., Lopes, I., Duarte, A. C., & Rocha-Santos, T. (2020). Environmental exposure to microplastics: An overview on possible human health effects. Science of the total environment, 702, 134455.

17. Leslie, H. A., Van Velzen, M. J., Brandsma, S. H., Vethaak, A. D., Garcia Vallejo, J. J., & Lamoree, M. H. (2022). Discovery and quantification of plastic particle pollution in human blood. Environment international, 163, 107199.

18. Horvatits, T., Tamminga, M., Liu, B., Sebode, M., Carambia, A., Fischer, L., … & Fischer, E. K. (2022). Microplastics detected in cirrhotic liver tissue. EBioMedicine, 82.

19. Yin, K., Wang, Y., Zhao, H., Wang, D., Guo, M., Mu, M., … & Xing, M. (2021). A comparative review of microplastics and nanoplastics: Toxicity hazards on digestive, reproductive and nervous system. Science of the total environment, 774, 145758.

20. Yuan, Z., Nag, R., & Cummins, E. (2022). Human health concerns regarding microplastics in the aquatic environment-From marine to food systems. Science of the Total Environment, 823, 153730.

21. Pan, D., Su, F., Liu, C., & Guo, Z. (2020). Research progress for plastic waste management and manufacture of value-added products. Advanced Composites and Hybrid Materials, 3, 443–461.

22. Singh, P., & Sharma, V. P. (2016). Integrated plastic waste management: environmental and improved health approaches. Procedia Environmental Sciences, 35, 692–700.

23. Shamsuyeva, M., & Endres, H. J. (2021). Plastics in the context of the circular economy and sustainable plastics recycling: Comprehensive review on research development, standardization and market. Composites Part C: open access, 6, 100168.

24. Soong, Y. H. V., Sobkowicz, M. J., & Xie, D. (2022). Recent advances in biological recycling of polyethylene terephthalate (PET) plastic wastes. Bioengineering, 9(3), 98.

25. Hopewell, J., Dvorak, R., & Kosior, E. (2009). Plastics recycling: challenges and opportunities. Philosophical Transactions of the Royal Society B: Biological Sciences, 364(1526), 2115–2126.

26. Ragaert, K., Delva, L., & Van Geem, K. (2017). Mechanical and chemical recycling of solid plastic waste. Waste management, 69, 24–58.

27. ISO. (2008). Plastics–Guidelines for the recovery and recycling of plastics waste. ISO 15270: 2008 (E).

28. Khalid, M. Y., Arif, Z. U., Ahmed, W., & Arshad, H. (2022). Recent trends in recycling and reusing techniques of different plastic polymers and their composite materials. Sustainable Materials and Technologies, 31, e00382.

29. Vollmer, I., Jenks, M. J., Roelands, M. C., White, R. J., Van Harmelen, T., De Wild, P., … & Weckhuysen, B. M. (2020). Beyond mechanical recycling: giving new life to plastic waste. Angewandte Chemie International Edition, 59(36), 15402–15423.

30. Suzuki, G., Uchida, N., Tanaka, K., Matsukami, H., Kunisue, T., Takahashi, S., … & Osako, M. (2022). Mechanical recycling of plastic waste as a point source of microplastic pollution. Environmental Pollution, 303, 119114.

31. Moharir, R. V., & Kumar, S. (2019). Challenges associated with plastic waste disposal and allied microbial routes for its effective degradation: a comprehensive review. Journal of Cleaner Production, 208, 65–76.

32. Woolnough, C. A., Yee, L. H., Charlton, T., & Foster, L. J. R. (2010). Environmental degradation and biofouling of ‘green’plastics including short and medium chain length polyhydroxyalkanoates. Polymer International, 59(5), 658–667.

33. Hu, L., He, L., Cai, L., Wang, Y., Wu, G., Zhang, D., … & Wang, Y. Z. (2024). Deterioration of single-use biodegradable plastics in high-humidity air and freshwaters over one year: Significant disparities in surface physicochemical characteristics and degradation rates. Journal of Hazardous Materials, 465, 133170.

34. Huo, Y., Dijkstra, F. A., Possell, M., & Singh, B. (2024). Mineralisation and priming effects of a biodegradable plastic mulch film in soils: Influence of soil type, temperature and plastic particle size. Soil Biology and Biochemistry, 189, 109257.

35. Zhang, Y., Cao, Y., Chen, B., Dong, G., Zhao, Y., & Zhang, B. (2024). Marine biodegradation of plastic films by Alcanivorax under various ambient temperatures: Bacterial enrichment, morphology alteration, and release of degradation products. Science of The Total Environment, 917, 170527.

36. Zhu, B., Wang, D., & Wei, N. (2022). Enzyme discovery and engineering for sustainable plastic recycling. Trends in biotechnology, 40(1), 22–37.

37. Park, S. Y., & Kim, C. G. (2019). Biodegradation of micro-polyethylene particles by bacterial colonization of a mixed microbial consortium isolated from a landfill site. Chemosphere, 222, 527–533.

38. Miri, S., Saini, R., Davoodi, S. M., Pulicharla, R., Brar, S. K., & Magdouli, S. (2022). Biodegradation of microplastics: Better late than never. Chemosphere, 286, 131670.

39. Tiwari, N., Santhiya, D., & Sharma, J. G. (2024). Significance of landfill microbial communities in biodegradation of polyethylene and nylon 6, 6 microplastics. Journal of Hazardous Materials, 462, 132786.

40. Cao, Z., Xia, W., Wu, S., Ma, J., Zhou, X., Qian, X., … & Jiang, M. (2023). Bioengineering Comamonas testosteroni CNB-1: a robust whole-cell biocatalyst for efficient PET microplastic degradation. Bioresources and Bioprocessing, 10(1), 94.

41. Taniguchi, I., Yoshida, S., Hiraga, K., Miyamoto, K., Kimura, Y., & Oda, K. (2019). Biodegradation of PET: current status and application aspects. Acs Catalysis, 9(5), 4089–4105.

42. Orlando, M., Molla, G., Castellani, P., Pirillo, V., Torretta, V., & Ferronato, N. (2023). Microbial enzyme biotechnology to reach plastic waste circularity: current status, problems and perspectives. International Journal of Molecular Sciences, 24(4), 3877.

43. Ahmed, S. F., Islam, N., Tasannum, N., Mehjabin, A., Momtahin, A., Chowdhury, A. A., … & Mofijur, M. (2024). Microplastic removal and management strategies for wastewater treatment plants. Chemosphere, 347, 140648.

44. Hettiarachchi, H., & Meegoda, J. N. (2023). Microplastic pollution prevention: the need for robust policy interventions to close the loopholes in current waste management practices. International journal of environmental research and public health, 20(14), 6434.

45. Ahmed, T., Shahid, M., Azeem, F., Rasul, I., Shah, A. A., Noman, M., … & Muhammad, S. (2018). Biodegradation of plastics: current scenario and future prospects for environmental safety. Environmental science and pollution research, 25, 7287–7298.

46. Chattopadhyay, I. (2022). Role of microbiome and biofilm in environmental plastic degradation. Biocatalysis and Agricultural Biotechnology, 39, 1022 63.

47. Koshti, R., Mehta, L., & Samarth, N. (2018). Biological recycling of polyethylene terephthalate: a mini-review. Journal of Polymers and the Environment, 26, 3520–3529.

48. Madhu, A., & Chakraborty, J. N. (2017). Developments in application of enzymes for textile processing. Journal of cleaner production, 145, 114–133.

49. Somarathna, H. M. C. C., Raman, S. N., Mohotti, D., Mutalib, A. A., & Badri, K. H. (2018). The use of polyurethane for structural and infrastructural engineering applications: A state-of-the-art review. Construction and Building Materials, 190, 995–1014.

50. Chattopadhyay, D. K., & Raju, K. V. S. N. (2007). Structural engineering of polyurethane coatings for high performance applications. Progress in polymer science, 32(3), 352–418.

51. Shi, J., Zheng, T., Zhang, Y., Guo, B., & Xu, J. (2021). Cross-linked polyurethane with dynamic phenol-carbamate bonds: properties affected by the chemical structure of isocyanate. Polymer Chemistry, 12(16), 2421–2432.

52. Akindoyo, J. O., Beg, M., Ghazali, S., Islam, M. R., Jeyaratnam, N., & Yuvaraj, A. R. (2016). Polyurethane types, synthesis and applications–a review. Rsc Advances, 6(115), 114453–114482.

53. Chattopadhyay, D. K., & Webster, D. C. (2009). Thermal stability and flame retardancy of polyurethanes. Progress in Polymer Science, 34(10), 1068–1133.

54. de Souza, F. M., Kahol, P. K., & Gupta, R. K. (2021). Introduction to polyurethane chemistry. In Polyurethane chemistry: Renewable polyols and isocyanates (pp. 1-24). American Chemical Society.

55. Christenson, E. M., Anderson, J. M., & Hiltner, A. (2007). Biodegradation mechanisms of polyurethane elastomers. Corrosion Engineering, Science and Technology, 42(4), 312–323.

56. Cregut, M., Bedas, M., Durand, M. J., & Thouand, G. (2013). New insights into polyurethane biodegradation and realistic prospects for the development of a sustainable waste recycling process. Biotechnology advances, 31(8), 1634–1647.

57. Bhavsar, P., Bhave, M., & Webb, H. K. (2023). Solving the plastic dilemma: the fungal and bacterial biodegradability of polyurethanes. World Journal of Microbiology and Biotechnology, 39(5), 122.

58. Pantelic, B., Skaro Bogojevic, S., Milivojevic, D., Ilic-Tomic, T., Lončarević, B., Beskoski, V., … & Nikodinovic-Runic, J. (2023). Set of small molecule polyurethane (PU) model substrates: Ecotoxicity evaluation and identification of PU degrading biocatalysts. Catalysts, 13(2), 278.

59. Sonnenschein, M. F., & Koonce, W. (2002). Polyurethanes. Encyclopedia of Polymer Science and Technology.

60. Zhang, J., & Hu, C. P. (2008). Synthesis, characterization and mechanical properties of polyester-based aliphatic polyurethane elastomers containing hyperbranched polyester segments. European Polymer Journal, 44(11), 3708–3714.

61. Fratesi, S. E., Lynch, F. L., Kirkland, B. L., & Brown, L. R. (2004). Effects of SEM preparation techniques on the appearance of bacteria and biofilms in the Carter Sandstone. Journal of Sedimentary Research, 74(6), 858–867.

62. Kim, H. W., Jo, J. H., Kim, Y. B., Le, T. K., Cho, C. W., Yun, C. H., … & Yeom, S. J. (2021). Biodegradation of polystyrene by bacteria from the soil in common environments. Journal of Hazardous Materials, 416, 126239.

63. Puiggené, Ò., Espinosa, M. J. C., Schlosser, D., Thies, S., Jehmlich, N., Kappelmeyer, U., … & Eberlein, C. (2022). Extracellular degradation of a polyurethane oligomer involving outer membrane vesicles and further insights on the degradation of 2, 4-diaminotoluene in Pseudomonas capeferrum TDA1. Scientific Reports, 12(1), 2666.

64. Gallego, V., García, M. T., & Ventosa, A. (2005). Methylobacterium hispanicum sp. nov. and Methylobacterium aquaticum sp. nov., isolated from drinking water. International Journal of Systematic and Evolutionary Microbiology, 55(1), 281–287.

65. Elahi, A., Bukhari, D. A., Shamim, S., & Rehman, A. (2021). Plastics degradation by microbes: A sustainable approach. Journal of King Saud University Science, 33(6), 101538.

66. Mahajan, N., & Gupta, P. (2015). New insights into the microbial degradation of polyurethanes. RSC Advances, 5(52), 41839–41854.

67. Huang, W. E., Hopper, D., Goodacre, R., Beckmann, M., Singer, A., & Draper, J. (2006). Rapid characterization of microbial biodegradation pathways by FT-IR spectroscopy. Journal of Microbiological Methods, 67(2), 273–280.

68. Dierkes, G., Lauschke, T., & Földi, C. (2021). Analytical methods for plastic (microplastic) determination in environmental samples. In Plastics in the Aquatic Environment-Part I: Current Status and Challenges (pp. 43–67). Cham: Springer International Publishing.

69. Spontón, M., Casis, N., Mazo, P., Raúd, B., Simonetta, A., Rios, L., & Estenoz, D. (2013). Biodegradation study by Pseudomonas sp. of flexible polyurethane foams derived from castor oil. International Biodeterioration & Biodegradation, 85, 85–94.

70. Fotopoulou, K. N., & Karapanagioti, H. K. (2015). Surface properties of beached plastics. Environmental Science and Pollution Research, 22, 11022–11032.

71. Romano, A., Rosato, A., Sisti, L., Zanaroli, G., Asadauskas, S. J., Nemaniutė, P., … & Totaro, G. (2024). Enzyme-catalyzed polyurethane adhesive degradation. Reaction Chemistry & Engineering.

72. Dias, R. C. M., Góes, A. M., Serakides, R., Ayres, E., & Oréfice, R. L. (2010). Porous biodegradable polyurethane nanocomposites: preparation, characterization, and biocompatibility tests. Materials Research, 13, 211–218.

73. Yang, X. F., Vang, C., Tallman, D. E., Bierwagen, G. P., Croll, S. G., & Rohlik, S. (2001). Weathering degradation of a polyurethane coating. Polymer degradation and stability, 74(2), 341–351.

74. Marchant, R. E., Zhao, Q., Anderson, J. M., & Hiltner, A. (1987). Degradation of a poly (ether urethane urea) elastomer: infra-red and XPS studies. Polymer, 28(12), 2032–2039.

75. Gupta, N., Rathi, P., & Gupta, R. (2002). Simplified para-nitrophenyl palmitate assay for lipases and esterases. Analytical biochemistry, 311(1), 98–99.

76. Castro-Ochoa, L. D., Rodríguez-Gómez, C., Valerio-Alfaro, G., & Ros, R. O. (2005). Screening, purification and characterization of the thermoalkalophilic lipase produced by Bacillus thermoleovorans CCR11. Enzyme and Microbial Technology, 37(6), 648–654.

77. Asoodeh, A., & Ghanbari, T. (2013). Characterization of an extracellular thermophilic alkaline esterase produced by Bacillus subtilis DR8806. Journal of Molecular Catalysis B: Enzymatic, 85, 49–55.

78. Howard, G. T. (2002). Biodegradation of polyurethane: a review. International Biodeterioration & Biodegradation, 49(4), 245–252.

79. Fuentes-Jaime, J., Vargas-Suárez, M., Cruz-Gómez, M. J., & Loza-Tavera, H. (2022). Concerted action of extracellular and cytoplasmic esterase and urethane-cleaving activities during Impranil biodegradation by Alicycliphilus denitrificans BQ1. Biodegradation, 33(4), 389–406.

80. Schmidt, J., Wei, R., Oeser, T., Dedavid e Silva, L. A., Breite, D., Schulze, A., & Zimmermann, W. (2017). Degradation of polyester polyurethane by bacterial polyester hydrolases. Polymers, 9(2), 65.

81. Jiang, W., Zhang, C., Gao, Q., Zhang, M., Qiu, J., Yan, X., & Hong, Q. (2020). Carbamate CN hydrolase gene ameH responsible for the detoxification step of methomyl degradation in Aminobacter aminovorans strain MDW-2. Applied and Environmental Microbiology, 87(1), e02005–20.

82. Vargas-Suárez, M., Fernández-Cruz, V., & Loza-Tavera, H. (2019). Biodegradation of polyacrylic and polyester polyurethane coatings by enriched microbial communities. Applied microbiology and biotechnology, 103, 3225–3236.

83. Katnic, S. P., de Souza, F. M., & Gupta, R. K. (2024). Recent Progress in Enzymatic Degradation and Recycling of Polyurethanes. Biochemical Engineering Journal, 109363.

84. Howard, G. T., & Blake, R. C. (1998). Growth of Pseudomonas fluorescens on a polyester–polyurethane and the purification and characterization of a polyurethanase–protease enzyme. International Biodeterioration & Biodegradation, 42(4), 213–220.

85. Oh, C., Kim, T. D., & Kim, K. K. (2019). Carboxylic ester hydrolases in bacteria: Active site, structure, function and application. Crystals, 9(11), 597.

86. Raczyńska, A., Góra, A., & André, I. (2024). An overview on polyurethane degrading enzymes. Biotechnology Advances, 108439.

87. Branson, Y., Söltl, S., Buchmann, C., Wei, R., Schaffert, L., Badenhorst, C.P.S., Reisky, L., Jäger, G., Bornscheuer, U.T., (2023). Urethanases for the enzymatic hydrolysis of low molecular weight carbamates and the recycling of polyurethanes. Angew. Chem. Int. Ed. Eng. 62 (9), e202216220.

88. Salgado, C. A., de Almeida, F. A., Barros, E., Baracat-Pereira, M. C., Baglinière, F., & Vanetti, M. C. D. (2021). Identification and characterization of a polyurethanase with lipase activity from Serratia liquefaciens isolated from cold raw cow’s milk. Food Chemistry, 337, 127954.

89. Masaki, K., Mizukure, T., Kakizono, D., Fujihara, K., Fujii, T., & Mukai, N. (2020). New urethanase from the yeast Candida parapsilosis. Journal of bioscience and bioengineering, 130(2), 115–120.

90. Sy, A., Timmers, A. C., Knief, C., & Vorholt, J. A. (2005). Methylotrophic metabolism is advantageous for Methylobacterium extorquens during colonization of Medicago truncatula under competitive conditions. Applied and Environmental Microbiology, 71(11), 7245–7252.

91. Dedysh, S. N., Knief, C., & Dunfield, P. F. (2005). Methylocella species are facultatively methanotrophic. Journal of bacteriology, 187(13), 4665–4670.

92. Yáñez, L., Rodríguez, Y., Scott, F., Vergara-Fernández, A., & Muñoz, R. (2022). Production of (R)-3-hydroxybutyric acid from methane by in vivo depolymerization of polyhydroxybutyrate in Methylocystis parvus OBBP. Bioresource technology, 353, 127141.

93. Pieja, A. J., Sundstrom, E. R., & Criddle, C. S. (2011). Poly-3-hydroxybutyrate metabolism in the type II methanotroph Methylocystis parvus OBBP. Applied and environmental microbiology, 77(17), 6012–6019.

94. Hiraishi, A., Furuhata, K., Matsumoto, A., Koike, K. A., Fukuyama, M., & Tabuchi, K. (1995). Phenotypic and genetic diversity of chlorine-resistant Methylobacterium strains isolated from various environments. Applied and Environmental Microbiology, 61(6), 2099–2107.

95. Jourand, P., Renier, A., Rapior, S., de Faria, S. M., Prin, Y., Galiana, A., … & Dreyfus, B. (2005). Role of methylotrophy during symbiosis between Methylobacterium nodulans and Crotalaria podocarpa. Molecular plant-microbe interactions, 18(10), 1061–1068.

96. Alamgir, K. M., Masuda, S., Fujitani, Y., Fukuda, F., & Tani, A. (2015). Production of ergothioneine by Methylobacterium species. Frontiers in Microbiology, 6, 1185.

97. Holland, M. A. (1997). Occam’s Razor Applied to Hormonology (Are Cytokinins Produced by Plants?). Plant Physiology, 115(3), 865.

98. Delmotte, N., Knief, C., Chaffron, S., Innerebner, G., Roschitzki, B., Schlapbach, R., … & Vorholt, J. A. (2009). Community proteogenomics reveals insights into the physiology of phyllosphere bacteria. Proceedings of the National Academy of Sciences, 106(38), 16428–16433.

99. Klikno, J., & Kutschera, U. (2017). Regulation of root development in Arabidopsis thaliana by phytohormone-secreting epiphytic methylobacteria. Protoplasma, 254(5), 1867–1877.

100. Mohanty, S. R., Mahawar, H., Bajpai, A., Dubey, G., Parmar, R., Atoliya, N., … & Kollah, B. (2023). Methylotroph bacteria and cellular metabolite carotenoid alleviate ultraviolet radiation-driven abiotic stress in plants. Frontiers in Microbiology, 13, 899268.

101. Tani, A., Takai, Y., Suzukawa, I., Akita, M., Murase, H., & Kimbara, K. (2012). Practical application of methanol-mediated mutualistic symbiosis between Methylobacterium species and a roof greening moss, Racomitrium japonicum. PLoS One, 7(3), e33800.

102. Tani, A., Ogura, Y., Hayashi, T., & Kimbara, K. (2015). Complete genome sequence of Methylobacterium aquaticum strain 22A, isolated from Racomitrium japonicum moss. Genome Announcements, 3(2), 10–1128.

103. Tsagkari, E., & Sloan, W. T. (2019). Impact of Methylobacterium in the drinking water microbiome on removal of trihalomethanes. International biodeterioration & biodegradation, 141, 10–16.

104. Calatrava, V., Hom, E. F., Llamas, Á., Fernández, E., & Galván, A. (2018). OK, thanks! A new mutualism between Chlamydomonas and methylobacteria facilitates growth on amino acids and peptides. FEMS microbiology letters, 365(7), fny021.

105. Ng, H. K. W. H. J., & Ivanova, E. P. (2014). 16 The Family Methylocystaceae.

106. Shah, Z., Hasan, F., Krumholz, L., Aktas, D. F., & Shah, A. A. (2013). Degradation of polyester polyurethane by newly isolated Pseudomonas aeruginosa strain MZA-85 and analysis of degradation products by GC–MS. International Biodeterioration & Biodegradation, 77, 114–122.

107. Nakajima-Kambe, T., Onuma, F., Akutsu, Y., & Nakahara, T. (1997). Determination of the polyester polyurethane breakdown products and distribution of the polyurethane degrading enzyme of Comamonas acidovorans strain TB-35. Journal of Fermentation and Bioengineering, 83(5), 456–460.

108. Zhang, Y., Xia, Z., Huang, H., & Chen, H. (2009). Thermal degradation of polyurethane based on IPDI. Journal of Analytical and Applied Pyrolysis, 84(1), 89–94.

109. Jiao, L., Xiao, H., Wang, Q., & Sun, J. (2013). Thermal degradation characteristics of rigid polyurethane foam and the volatile products analysis with TG-FTIR-MS. Polymer Degradation and Stability, 98(12), 2687–2696.

110. do Canto, V. P., Thompson, C. E., & Netz, P. A. (2019). Polyurethanases: Three-dimensional structures and molecular dynamics simulations of enzymes that degrade polyurethane. Journal of Molecular Graphics and Modelling, 89, 82–95.

